# Emerging Concern of Scientific Fraud: Deep Learning and Image Manipulation

**DOI:** 10.1101/2020.11.24.395319

**Authors:** Chang Qi, Jian Zhang, Peng Luo

## Abstract

Scientific fraud by image duplications and manipulations within western blot images is a rising problem. Currently, problematic western blot images are mainly detected by checking repeated bands or through visual observation. However, the completeness of the above methods in detecting problematic images has not been demonstrated. Here we show that Generative Adversarial Nets (GANs) can generate realistic western blot images that indistinguishable from real western blots. The overall accuracy of researchers for identifying synthetic western blot images is 0.52, which almost equal to blind guess (0.5). We found that GANs can generate western blot images with bands of the expected lengths, widths, and angles in desired positions that can fool researchers. For the case study, we find that the accuracy of detecting the synthetic western blot images is related to years of researchers performed studies relevant to western blots, but there was no apparent difference in accuracy among researchers with different academic degrees. Our results demonstrate that GANs can generate fake western blot images to fool existing problematic image detection methods. Therefore, more information is needed to ensure that the western blots appearing in scientific articles are real. We argue to require every western blot image to be uploaded along with a unique identifier generated by the laboratory machine and to peer review these images along with the corresponding submitted articles, which may reduce the incidence of scientific fraud.

---

Western blotting is an analytical technique widely used in molecular biology and immunogenetics research^1^. However, since the 1990s, the accuracy and reproducibility of the western blot-based analysis of proteins has begun to be questioned by the academic community. In addition, many of the problematic images found in scientific papers are related to western blots^2^. Bik E M. et al. scanned a total of 20621 papers published from 1995 to 2014. The results of their study suggest that one of every 25 articles that contain western blot images or other images may suffer from data anomalies^3^. On the well-known academic antifraud platform “Retraction Watch” (www.retractionwatch.com), the keyword “western blot” brings up over 200 reports. Currently, problematic western blot images are mainly detected by checking repeated bands or through visual observation^3^. However, the above methods can detect only problematic images that have been manipulated based on existing western blot images, while generated western blot images not based on publicly available western blots are missed.

Recently, there has been great progress in the field of generative adversarial networks (GANs)^4^ and their extensions. A GAN operates on the basis of a minimax game between two neural networks: the generator generates samples, and the discriminator identifies the sources of samples. In this game, the adversarial loss incurred by the discriminator provides a clever way to capture high-dimensional and complex distributions by imposing higher-order consistency, which has been proven to be useful for generating realistic images for various applications in the computer vision field, such as plausible image generation^5^, text-to-image generation^6^, image-to-image translation^7^, image completion^8^, and super-resolution^9^. The diverse and successful applications of GANs in image generation tasks have attracted growing interest across many communities. To investigate whether GANs can generate realistic western blot images, we trained several GAN models to fit a western blot distribution and generated western blots using the trained models.

We collected western blots from the laboratory, exported them as images in .tif format and canceled the invert parameter in the exported .tif images. Then, we performed data preprocessing, as shown in Fig 1A, on the collected western blot images to generate corresponding semantic images. First, the western blot images were binarized to generate images with pixel values of only 0 and 1, where 0 represents background and 1 represents a band region (foreground). Second, each band’s edge was outlined using the ‘findcontour’ method in OpenCV^10^, and then, the ‘minAreaRect’ function was used to obtain each edge’s minimum outer rectangle. We established several rules for screening out undesirable western blots, which can be grouped into five categories^11^: (1) unusual or unexpected bands, (2) no bands, (3) faint bands or weak signal, (4) high background on the blot, and (5) patchy or uneven spots on the blot. Finally, we obtained a dataset containing 233 pairs of western blots and their corresponding semantic images representing the position, length, width, and angle of each band.

**Fig. 1.**
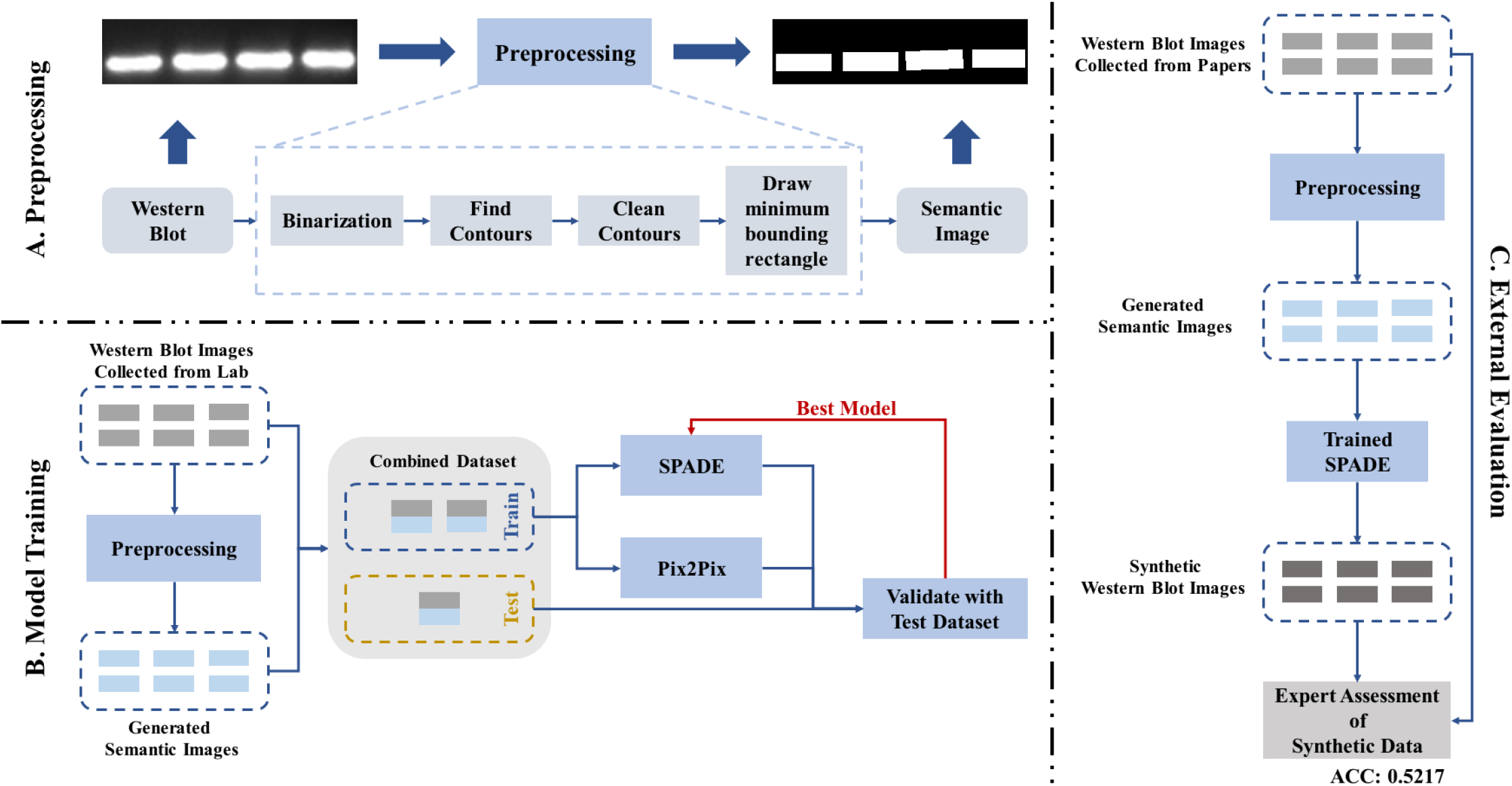
This is a Flowchart of the proposed method. A. Data preprocessing to obtain paired datasets. B. Model training procedure. C. External evaluation process.

As shown in Fig 1B, we divided the paired dataset into a training set and a test set. The test set contained 30 pairs, and the training set contained 203 pairs. We first tried to use CycleGAN^7^ to model the training set by randomly generating semantic-like images to teach the generator the general style to be translated, rather than obtaining corresponding western blot and semantic image pairs following the preprocessing process described above. However, the experimental results showed that consistency of the paired images is greatly needed for the translation process. Therefore, we chose to use the paired image-to-image translation models pix2pix^12^ and SPADE^13^ to model the training set. Specifically, each model includes a generator G, which aims to generate western blot images based on input semantic images. Each model also includes a discriminator D, which attempts to recognize real western blot images and synthetic western blot images. The purpose when training D is to allow it to recognize synthetic images as accurately possible. Meanwhile, the purpose when training G is to allow it to fool D as effectively as possible, i.e., to produce candidates that the trained discriminator will classify as real western blots (i.e., as part of the true data distribution). For training, backpropagation is applied to both the generator and the discriminator alternately so that they are both forced to make better decisions. The generator is forced to produce images that fit the true data distribution, and the discriminator becomes more skilled at detecting synthetic western blot images. In SPADE, a spatially adaptive normalization layer is used to inject the input semantic image into the generator, with the aim of better preserving semantic information compared with the case in which the semantic image is fed to the generator directly (pix2pix).

To validate the capabilities of pix2pix and SPADE in modeling western blot images based on given semantic images, the semantic images from the test set were input into the trained generators, and the synthetic western blots produced by the two generators were compared to the real western blots. The mean square error (MSE) matrix was calculated to represent the differences between the real observations and the observations predicted by each model. Since the MSE matrix for SPADE was better than that for pix2pix, indicating that SPADE could better model the real western blot image distribution, we chose SPADE to perform the subsequent external evaluation, as shown in Fig 1C.

For the external evaluation, we first collected western blot images from published articles and applied the same preprocessing process to obtain corresponding semantic images. In this way, we collected a total of 400 pairs of western blots and semantic images. Similar to the internal evaluation process, we then fed the semantic images into the SPADE generator trained on the data collected from the laboratory and obtained synthetic western blots. Subsequently, we published a questionnaire on the Internet and invited researchers in fields relevant to western blotting to distinguish the synthetic western blots from the real western blots. Our system randomly selected 40 of the 400 paired real and synthetic western blots for each researcher to classify. In addition, we recorded some basic information about the researchers participating in the study, such as years of relevant research and academic degrees. We analyzed the accuracy based on each participant’s basic information and conducted a case study based on the accuracy for each real and synthetic western blot pair.

We collected a total of 405 valid answers to the questionnaire to conduct relevant analyses. Fig 2A and Fig 2B show the distributions of the participants grouped by years of relevant research and academic degree. First, the overall accuracy among all participants was 0.52, indicating that the synthetic western blots were virtually indistinguishable from the real western blots; these results prove that it is possible for GANs to generate western blot images with bands of the expected lengths, widths, and angles in desired positions that are able to fool researchers. Second, researchers who had performed studies relevant to western blots over the past 5-10 years achieved the highest accuracy in detecting synthetic western blots (0.5831), whereas researchers who had performed such studies for less than 3 years and more than 10 years showed no noticeable difference in accuracy (0.4984 and 0.4975), and researchers who had performed studies for between 3 and 5 years achieved a classification accuracy of 0.5540. There was no apparent difference in accuracy among researchers with different academic degrees (0.5361 for M.D., 0.5373 for Ph.D., 0.5296 for M.M., and 0.5357 for B.M.).

**Fig. 2.**
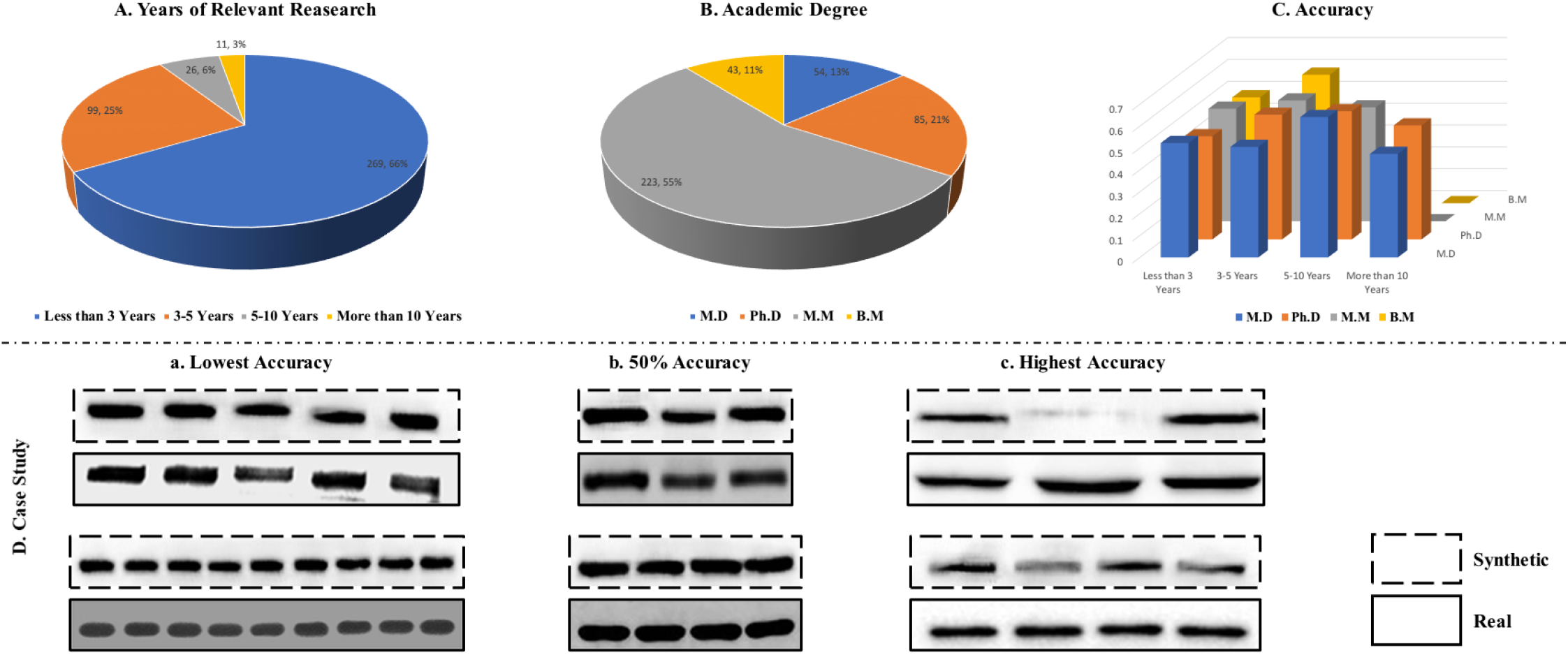
Results of External Evaluation.

Then, we calculated the overall accuracy for each synthetic and real western blot pair and separated the pairs into 3 groups with different levels of accuracy: the lowest accuracy (only 4/23 | 9/37 participants detected the synthetic western blot), 50% accuracy (the participants had difficulty distinguishing the synthetic western blot from the real one), and the highest accuracy (20/24 | 26/32 participants made the correct choice). In the group of western blots with the lowest accuracy, the synthetic images showed more noise in the background, consistent with the distribution of the real western blots, and the real western blots exhibited unusual patterns in band shape, which may have misled the participants’ choices. In the 50% accuracy group, the synthetic and real western blots exhibited the same patterns, noisy backgrounds, and regular band shapes, making them difficult for the participants to distinguish between. For the blots with the highest accuracy, some bands in the synthetic blots did not appear sufficiently realistic, which helped the participants identify the fake images.

There are some limitations to this work. First, we used a one-to-one image translation model, limiting the model’s ability to generate western blots. There may be many western blots in the real world that follow the same semantic pattern, with the same sizes, positions, and angles for the bands. However, in our model, a given semantic image can be used to generate only one western blot image. Second, we can control only the positions, angles, and sizes of the bands in the generated blots, which is another limitation of the model’s generation capabilities. In future work, we can attempt to generate blots with more controllable parameters, such as the shapes and gray values of the bands and the type of background noise. We will also try to achieve one-to-many translation by adding noise during the translation process and modifying the model structure to make it more controllable.

In summary, it is possible to generate fake western blot images by means of GANs; therefore, more information is needed to ensure that the western blots appearing in scientific articles are real. An effective solution might be to require every western blot image to be uploaded along with a unique identifier generated by the laboratory machine and to peer review these images along with the corresponding submitted articles, which may reduce the incidence of scientific fraud.

